# Bloodstream-associated *Salmonella* Typhimurium and Enteritidis iNTS pathovariants hyper-replicate in human macrophages

**DOI:** 10.64898/2026.01.28.702237

**Authors:** Francesco Amadeo, Marie Held, Shaun H. Pennington, Giancarlo A. Biagini, Jay C.D. Hinton, Natalia Cattelan

## Abstract

Invasive non-typhoidal *Salmonella* (iNTS) are a major cause of bloodstream infections in sub-Saharan Africa, yet the host–pathogen interaction mechanisms remain poorly understood. Here, we developed and optimised a human macrophage infection model based on PMA-differentiated THP-1 cells to investigate infection dynamics of clinically relevant *Salmonella* Typhimurium and *Salmonella* Enteritidis strains. We compared intracellular survival and replication of gastroenteritis-associated and bloodstream-associated pathovariants, including *S*. Typhimurium ST313 Lineage 2, the novel *S*. Typhimurium ST313 Lineage 3 and the understudied *S*. Enteritidis Central/Eastern African (CEAC) clades. Our results reveal that the CEAC *S*. Enteritidis and ST313 *S*. Typhimurium iNTS pathovariants hyper-replicate within host cells, compared to global epidemic isolates. The cellular model achieved robust pro-inflammatory polarisation of human macrophages, while revealing limitations in modelling macrophage plasticity. Overall, this work defines clear differences in intracellular behaviour of *Salmonella* pathovariants and establishes a robust experimental framework for future studies on invasive disease pathogenesis and therapeutic interventions that target iNTS bacteria.

## INTRODUCTION

Invasive non-typhoidal *Salmonella* (iNTS) infections are a leading cause of bloodstream infection in sub-Saharan Africa (sSA), with an estimated 535,000 cases and over 77,500 deaths worldwide in 2017 ^1,2^. Individuals at highest risk in low-income settings include young children affected by malaria, malnutrition or human immunodeficiency virus (HIV), as well as HIV-positive adults ^1-3^. In sSA, approximately two-thirds of iNTS infections are caused by *S*. Typhimurium, with most cases associated with the ST313 multilocus sequence type ^4-8^. ST313 is phylogenetically distinct from the globally dominant ST19 clade, which typically causes self-limiting gastroenteritis ^8^. *S*. Enteritidis causes approximately one-third of iNTS cases in sSA, with two monophyletic clades, the Central-Eastern African clade (CEAC) and Western African clade (WAC) ^7^.

In Africa, human iNTS disease results from systemic infection by *S*. Enteritidis or *S*. Typhimurium that is associated with the intracellular reticulo-endothelial niche, involving both blood-borne and bone-marrow monocytic cells ^9^. Following ingestion of contaminated food or water and colonisation of the gastrointestinal tract, *Salmonella* can invade intestinal epithelial cells and then be internalised by local dendritic cells and macrophages ^10^.

Macrophages are key components of the innate immune system; although designed to kill bacteria, macrophages serve as the primary intracellular niche for *Salmonella*. Systemic dissemination of non-typhoidal *Salmonella* involves macrophage-mediated transport to extra-intestinal sites such as liver and spleen ^11^. Within macrophages, *Salmonella* uses well-characterised type III secretion systems (T3SS) to translocate effector proteins to modulate the phagosome maturation process and generate the *Salmonella*-containing vacuole (SCV) that enables intracellular survival and replication ^12-14^. Infected macrophages subsequently migrate to mesenteric lymph nodes, facilitating systemic spread ^15^. We hypothesise that the *Salmonella* lineages that have evolved in the African setting will either possess enhanced mechanisms for counteracting macrophage host-defence factors or suppress intra-macrophage killing by subverting host signalling pathways.

Previously, a range of murine, bovine and avian animal infection models have been used to study the virulence of a range of *Salmonella* serovars ^16^. However, animal models have not successfully recapitulated the key phenotypes of iNTS disease seen in humans ^17,18^. Most research on intra-macrophage *Salmonella* infection has focused on murine systems, which have limited relevance for human-adapted iNTS bacteria. Previous studies using donor-derived primary human macrophages (Le Bury et al., 2021) suggested that *S*. Typhimurium ST313 replicated more efficiently than *S*. Typhimurium ST19, However, donor-to-donor variability between the primary macrophages meant that these findings were not statistically significant. The limitations of existing infection approaches prompted us to develop a human macrophage-derived cell model for the study of iNTS *Salmonella* infections.

Macrophage plasticity, the remarkable ability of these cells to adapt to exogenous signals like cytokines and PAMPs, gives rise to distinct polarisation states with unique specialised roles ^19,20^. Although such diverse states cannot be fully replicated *in vitro*, reproducible models involving macrophage-like cells are crucial for investigating host-pathogen interactions. The human monocytic leukaemia THP-1 cell line is widely used as a surrogate for primary macrophages ^21-23^, and has a track record of being used for *Salmonella* infection studies ^24-27^. THP-1 cells can be differentiated using phorbol 12-myristate-13-acetate (PMA) ^28-30^. Carefully optimised differentiation protocols, such as those established by Steele-Mortimer and colleagues, have provided robust and biologically informative macrophage models for *Salmonella* infection studies ^24^. However, substantial methodological discrepancies persist across the field, with PMA concentration, exposure time, and resting conditions varying widely between studies (Supplementary Table 1), highlighting the lack of a standardised differentiation protocol.

Here, we optimised a differentiation protocol for generating macrophage-like cells from THP-1 monocytes. The resulting *in vitro* model was successfully used to study infection dynamics and intracellular survival of three representative *S*. Typhimurium and four *S*. Enteritidis clinical isolates.

Our study reveals that iNTS Enteritidis and Typhimurium display a shared hyper-replication phenotype within human macrophages. *S*. Enteritidis CEAC strains D7795 and CP255 exhibit markedly greater intracellular survival and replication than the global epidemic clade (GEC) *S*. Enteritidis strains P125109 and A1636. Similarly, *S*. Typhimurium ST313 Lineage 2 (L2) and Lineage 3 (L3) show significantly enhanced intracellular survival compared with the gastroenteritis-associated ST19 clade.

## RESULTS

### PMA induces differentiation of THP-1-derived macrophages without triggering activation

To establish a human macrophage infection model suitable for studying intracellular infection by *S*. Typhimurium and *S*. Enteritidis, we first optimised a protocol to differentiate THP-1 monocytes using phorbol 12-myristate 13-acetate (PMA). Differentiation success was assessed by evaluation of cell morphology, adherence, and the expression of key surface markers across several PMA treatment conditions (Figure 1). Phase contrast imaging showed increased cell adherence under all four tested conditions (Figure 1A). While treatment with 10 ng/mL PMA for 24 h (Condition A) produced cells with a round, heterogeneous morphology, incubation with 50 ng/mL PMA for 48 h (Condition D) generated flattened, homogenous cells (Figure 1A). Flow cytometry revealed that the THP-1 cells exhibited heterogeneous expression of CD11b, CD14, and CD36 (Figure 1B). Condition A led to upregulation of CD14 and CD36 but no increase in CD11b levels (Figure 1B). In contrast, extended PMA treatment (Conditions B–D) yielded a uniform CD11b+, CD14+, and CD36+ population (Figure 1B), a finding confirmed by median fluorescence intensity analysis (Figure 1C).

**Figure 1.**
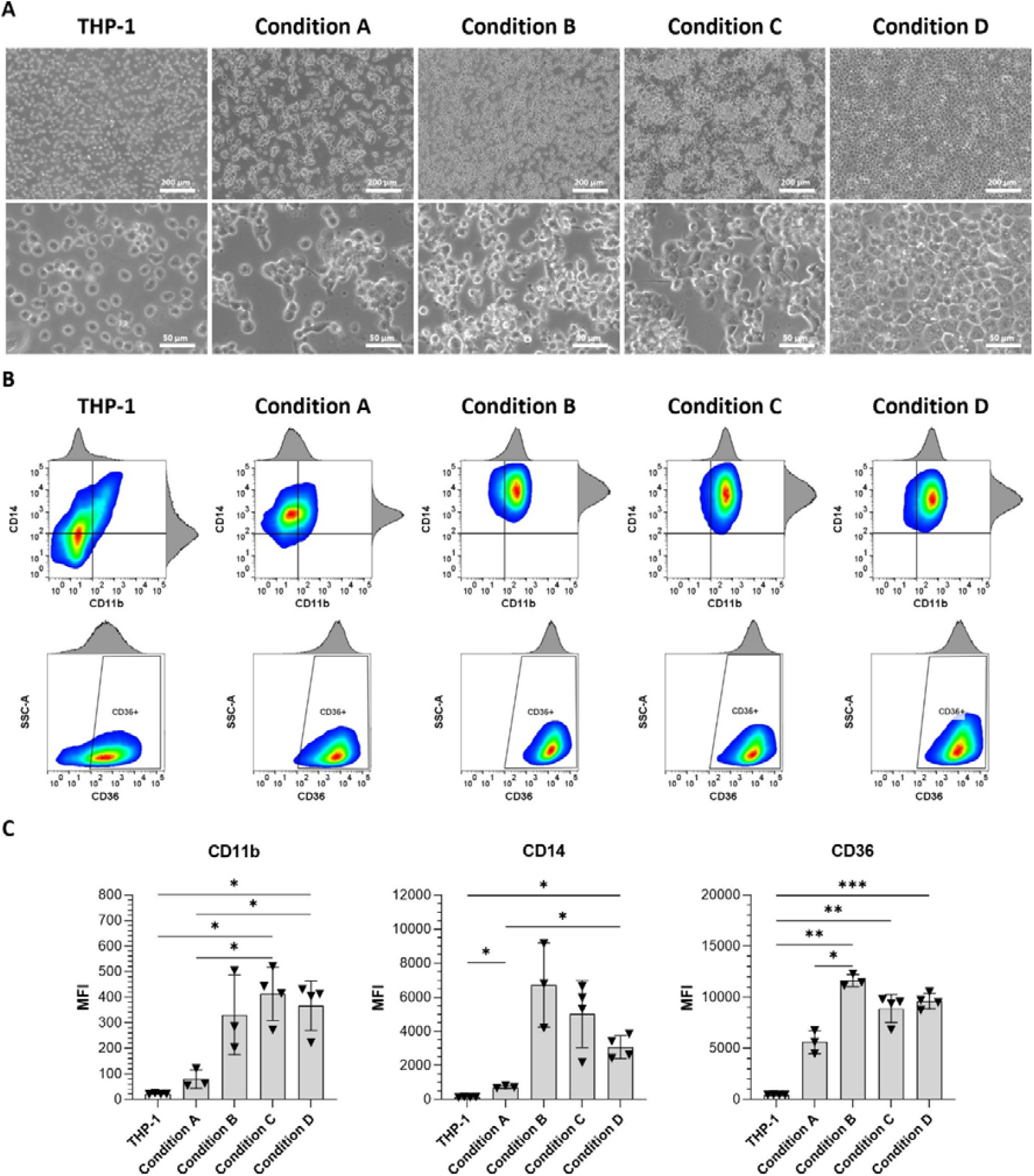
Cell morphology and macrophage surface markers are affected by PMA treatment. (A) Representative phase contrast images of THP-1 cells untreated versus THP-1 cells differentiated with Condition A (PMA 10 ng/mL for 24 h), Condition B (PMA 10 ng/mL for 24 h followed by a 24 h recovery in fresh media), Condition C (PMA 10 ng/mL for 48 h) and Condition D (PMA 50 ng/mL for 48 h). (B) Representative flow cytometric analysis of THP-1 cells and macrophage-induced cells stained using anti-CD11b, anti-CD14 and anti-CD36 antibodies. (C) Median fluorescence intensity (MFI) of CD11b, CD14 and CD36. Data are displayed as mean ± SD from n ≥ 3. Statistical analyses performed using the Brown-Forsythe and Welch ANOVA test and the Dunnet T3 multiple comparison post hoc test. * p < 0.05, ** p < 0.01, *** p < 0.001.

To verify that PMA treatment did not activate macrophages, we evaluated the surface expression of M1 and M2 markers by flow cytometry. None of the protocols tested caused biologically significant increases in CD86 and HLA-DR (M1) or CD163 and CD206 (M2), indicating that the PMA-induced differentiation did not stimulate macrophage activation (Supplementary Figure 1).

### THP-1-derived macrophages undergo M1 but not M2 polarisation

Macrophages are versatile immune cells capable of assuming a range of functional states. To evaluate the capacity of THP-1-derived macrophages to polarise into M1 (pro-inflammatory) or M2 (anti-inflammatory) states, cells generated by extended PMA treatment (Conditions B, C, and D) were treated with LPS and IFN-γ for 16 h or treated with IL-4 and IL-13 for 72 h, following established protocols for macrophage polarization. Condition A was excluded due to inadequate CD11b marker expression (Figure 1B).

All three protocols induced M1 polarisation, as indicated by elevated HLA-DR and CD86 surface expression (Supplementary Figure 2A-B), upregulation of IL-1β, TNFα, and HLA-DR mRNA (Supplementary Figure 2C), and increased secretion of IL-6 and TNFα (Supplementary Figure 2D). In contrast, none of the treatments induced a full M2 phenotype; CD163 and CD206 levels remained unchanged following IL-4/IL-13 stimulation (Supplementary Figure 2A-B), while only CCL-18 and CCL-22 mRNA were slightly upregulated. No IL-10 or TGFβ1 secretion was detected (Supplementary Figure 2D). Instead, the three treatments resulted in partial activation, indicated by increased HLA-DR and CD86 surface expression and HLA-DR mRNA levels (Supplementary Figure 2).

### THP-1-derived macrophages as a model for infection with Salmonella iNTS isolates

We identified three PMA-based protocols (Conditions B, C, and D) that generated macrophage-like cells from THP-1 monocytes. Due to increased cell detachment under Conditions B and C, Condition D (50 ng/mL PMA for 48 h) was selected for subsequent infection studies. Using this protocol, we assessed the intracellular survival of three *S*. Typhimurium and four *S*. Enteritidis clinical isolates, representing both global (4/74, P125109, A1636) and epidemic lineages (ST313 L2 D23580, ST313 L3 BKQZM9, CEAC D7795, CEAC CP255).

To optimise infection conditions, we compared multiplicities of infection (MOI) of 5 and 10. Colony-forming unit (CFU) assays revealed successful infection and replication by all strains across 24 h, regardless of MOI (Supplementary Figure 3A-B). MOI 10 resulted in higher bacterial counts at all time points (1.5, 4, 8, 24 hpi), though normalised survival curves were comparable at both MOIs (Supplementary Figure 3A-C). The MOI of 10 was used for all subsequent experiments (Figure 2).

**Figure 2.**
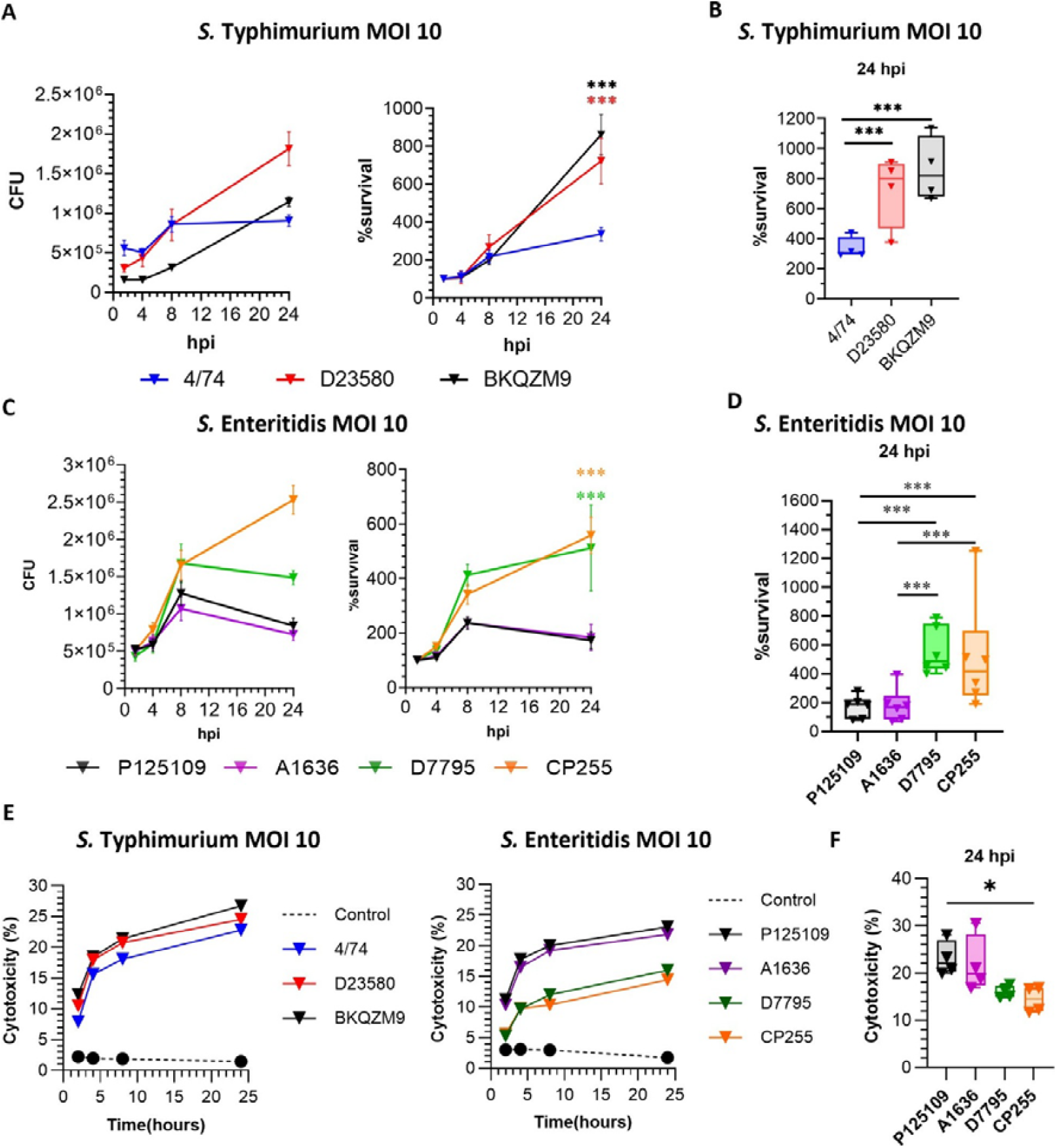
Bloodstream associated *Salmonella* strains survive better than gastroenteritis associated strains in THP-1 derived macrophages. THP-1 cells infected at a MOI of 10 with selected (A-B-E) *S*. Typhimurium ST19 (4/74) and ST313 L2 (D23580) and L3 (BKQZM9) isolates and (C-D-F) *S*. Enteritidis GEC (P125109 and A1636) and CEAC (D7795 and CP255) isolates. (A-D) Intracellular bacteria were enumerated via CFU at 1.5, 4, 8 and 24 hpi. Percent survival was expressed as the number of intracellular bacteria at each time point divided by the number of intracellular bacteria at 1.5 hpi and expressed as percentage. Data are displayed as mean ± SD from n = 6. Statistical analyses performed using the Brown-Forsythe and Welch ANOVA test and the Dunnet T3 multiple comparison post hoc test. * p < 0.05, ** p < 0.01. **(E-F)** Cytotoxicity (%) induced by *S*. Typhimurium (E) and *S*. Enteritidis (F) over 24 hours from infection compared to control not infected cells. Data are displayed as mean, no SD included, from n = 5.

All *S*. Typhimurium strains tested replicated intracellularly in human macrophages, with ST313 L3 BKQZM9 showing reduced initial uptake but enhanced proliferation by 24 hpi (Figure 2A). All strains followed a similar infection dynamic, with a minimal increase during the first 4 hpi, followed by active replication between 4 and 24 hpi. By 24 hpi, the ST313 L2 and L3 strains showed dramatically higher replication than the global ST19 strain 4/74 (Figure 2B).

The *S*. Enteritidis strains displayed comparable initial infection levels but different levels of intracellular replication. While global strains (P125109, A1636) showed restricted survival between 8 and 24 hpi, the CEAC strains (D7795, CP255) continued to proliferate (Figure 2C-D), resulting in significantly higher levels of intracellular replication at 24 hpi.

### Assessment of Bacterial Cytotoxicity and Single-Cell Replication Patterns

To determine the ability of the different pathovariants to kill human macrophages, cytotoxicity was evaluated with SYTOX Green staining (Figure 2E-F). Early cytotoxicity levels were consistently low for all *S*. Typhimurium strains (8.7–11.8%). By 24 hpi, cell death had increased slightly to ∼24–27%. The *S*. Enteritidis strains also caused low cytotoxicity initially, with CEAC strains D7795 and CP255 causing less cell death at 24 hpi (∼15–17%) than the global strains (∼22–23%), although only the difference between CP255 and P125109 was statistically significant (p<0.05). Cytotoxicity levels were not influenced by the two different MOI conditions. (Supplementary Figure 3D).

To investigate intracellular replication at the single-cell level, confocal microscopy was used to analyse infected THP-1 cells at multiple time points (Figure 3A). Infection rates were similar among *S*. Enteritidis isolates, while ST313 L3 BKQZM9 infected fewer cells than D23580 and 4/74, consistent with CFU results (Figure 3B). Intracellular bacterial counts increased over time for all strains (Figure 3C). Quantification of the number of intracellular bacteria showed no differences at early times post infection between the *S*. Typhimurium isolates (correlating again with our CFU data). In contrast, *S*. Enteritidis CEAC isolates produced a rapid increase in bacterial burden, with significant differences in the number of intracellular bacteria at 4 and 8 hpi, involving some clusters of over 20 bacteria per cell (Figure 3C).

**Figure 3.**
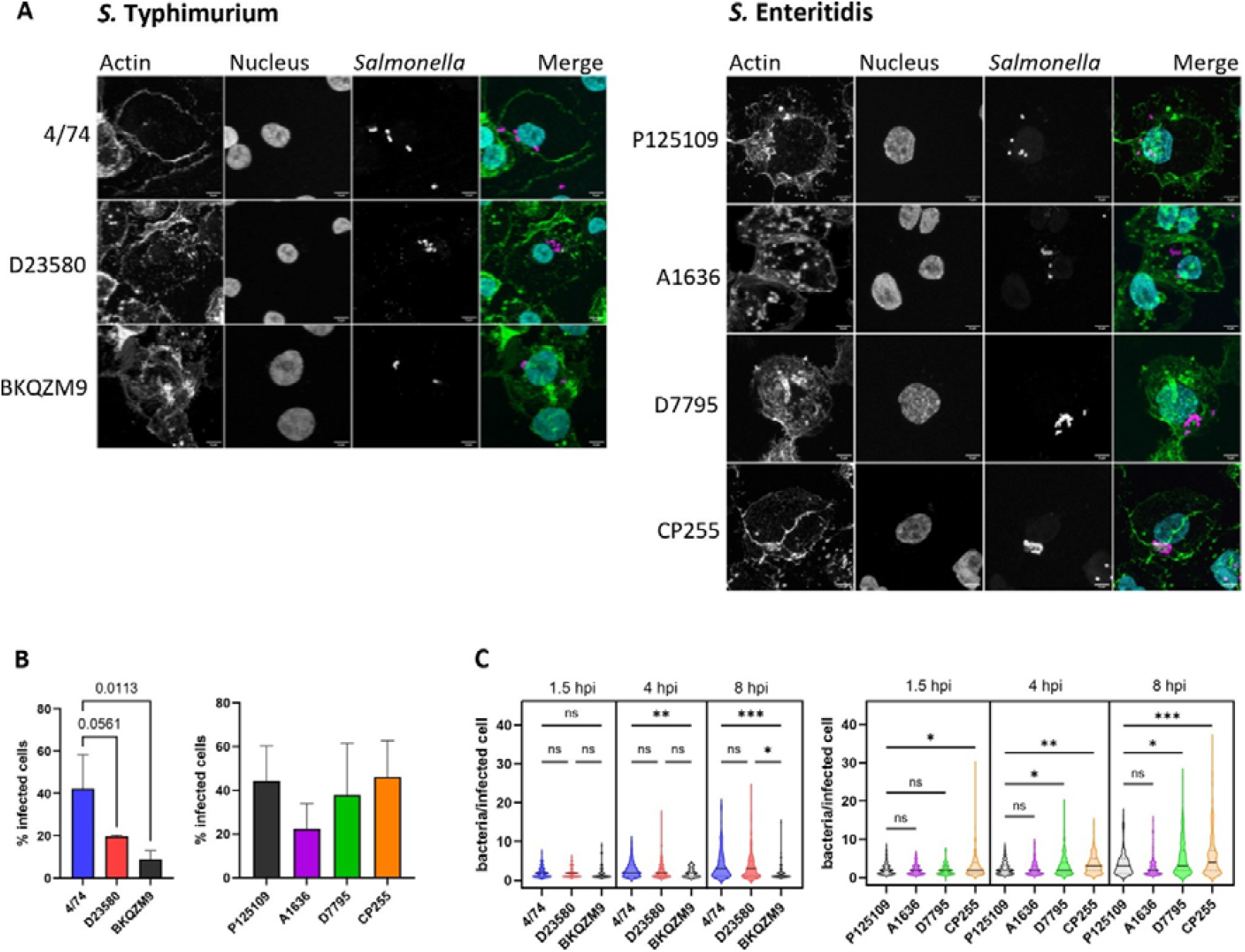
Visualization of THP-1 derived macrophages infected with *Salmonella*. THP-1 cells infected at an MOI of 10 with selected (A) *S*. Typhimurium ST19 (4/74) and ST313 L2 (D23580) and L3 (BKQZM9) isolates (left panels) and *S*. Enteritidis GEC (P125109 and A1636) and CEAC (D7795 and CP255) isolates (right panels). Cells were fixed and stained for F-actin (green), LPS (magenta) and label DNA (cyan). Representative images of three independent experiments at 8 hpi are shown. (B) Image analysis was performed with Cellpose and StarDist. Percentage of infected cells deduced from the number of cells containing intracellular bacteria at 1.5 hpi divided by the total number of cells. (C) The number of intracellular bacteria per cell was scored at 1.5, 4 and 8 hpi. Data correspond to three independent experiments. Data are displayed as mean ± SD from n = 3. Statistical analyses performed using the Brown-Forsythe and Welch ANOVA test and the Dunnet T3 multiple comparison post hoc test. * p < 0.05, ** p < 0.01, *** p<0.001.

## DISCUSSION

The ability of invasive non-typhoidal *Salmonella* (iNTS) to persist and replicate within human macrophages is likely to be a key strategy for avoiding detection and clearance by the host immune system. Therefore, development of a robust human macrophage infection model is critical for investigating host–pathogen interactions, elucidating bacterial survival mechanisms, and identifying features that differentiate iNTS from gastroenteritis-associated strains.

Here, we established a THP-1-derived macrophage model suitable for studying infection and intracellular survival of representative *S*. Typhimurium and *S*. Enteritidis pathovariants. We have used the standardised THP-1 model to do the first direct comparison of a gastroenteritis-associated *S*. Typhimurium ST19 isolate with ST313 Lineages 2 and 3, and between *S*. Enteritidis strains from both the GEC (P125109 and A1636) and the CEAC (D7795 and CP255) clades.

Existing methods for inducing differentiation of THP-1 cells into cells that resemble monocyte-derived macrophages (MDMs) rely on PMA treatment ^24,29,30^. However, published protocols involved a wide variety of PMA concentrations, incubation times, and recovery periods (Supplementary Table 1). Since no universal, standardised protocol existed for differentiating THP-1 cells ^31^ we optimised the method by comparing low (10 ng/mL) and moderate (50 ng/mL) PMA concentrations. Because higher PMA doses and longer exposures can activate macrophages, concentrations above 50 ng/mL were avoided ^24,32,33^.

We found that protocols involving treatment with 10 ng/mL PMA for 24 h with a 24 h recovery (Condition B), 48 h treatment (Condition C), or 50 ng/mL for 48 h (Condition D) yielded macrophage-like phenotypes with no activation ^34^.

Individuals in many low-income countries who are immunocompromised, such as HIV-positive adults or children affected by malaria or malnutrition, are especially vulnerable to iNTS infections ^3^ and often exhibit an M2-skewed macrophage phenotype ^35^. Replication of *S*. Typhimurium ST19 in human MDMs is known to be phenotype-dependent: M1 macrophages inhibit bacterial growth, whereas M2a and M0 cells permit replication ^36^. Because *S*. Typhimurium can drive infected macrophages towards a replication-permissive M2-like state ^37 38^, we tested the abilities of our THP-1-derived macrophages to differentiate into specific phenotypes. Treatment with LPS and IFN-γ successfully induced M1 polarisation in Conditions B, C, and D, as has been reported previously ^34,39^. Conversely, IL-4 and IL-13 treatment for 72 h did not generate M2-like phenotypes, consistent with previous findings on the variability of THP-1 responses ^28^. Accordingly, Condition D was adopted as our optimised THP-1 differentiation protocol. The findings highlight the need for standardised protocols and careful interpretation when using THP-1 models.

We next used our optimised model to examine macrophage infection with a range of non-typhoidal *Salmonella* strains. Our panel included three *S*. Typhimurium strains (ST19 4/74; ST313 L2 D23580; ST313 L3 BKQZM9) and four *S*. Enteritidis strains (GEC P125109, GEC A1636; CEAC D7795, CEAC CP255). Once engulfed by host macrophages, *Salmonella* bacteria experience a range of fates ranging from persistence, replication, the induction of cell death or intracellular destruction ^40,41^. All tested strains successfully infected and replicated in macrophages for up to 8 hpi. Our findings for *S*. Typhimurium ST313 L2 align with earlier studies ^25,26^. However, we observed higher intracellular survival of *S. Typhimurium* ST19 than previous reports. Whereas Ramachandran et al. reported ∼50% survival at 24 hpi, we observed ∼290% survival under our conditions. This difference likely reflects variation between THP-1 differentiation protocols and bacterial isolates, as we used strain 4/74 while Ramachandran et al. used strains I77 and SL1344 ^26^.

We report for the first time the intra-macrophage infection and replication capacity of S. Typhimurium ST313 L3 ^8^. Despite lower uptake, this strain showed enhanced replication over 24 h. ST313 L3 is a phylogenetic intermediate between ST313 L1 and L2 that shares 99.98% nucleotide sequence identity with D23580 ^8^. ST313 L3 also had increased virulence in a murine infection model ^42^ and a higher genome-derived invasiveness index than ST313 L2 ^8^. Clearly, the virulence mechanisms of ST313 L3 warrant further investigation.

The CEAC and WAC lineages of *S*. Enteritidis cause about one-third of iNTS cases in sub-Saharan Africa ^7,43,44^. Until now, the intracellular behaviour of the CEAC lineage had only been studied in murine RAW264.7 cells ^45^. Here we show that GEC (P125109, A1636) and CEAC (D7795, CP255) isolates were internalised in THP-1 cells at similar levels, but the CEAC isolates exhibited significantly greater replication between 8 hpi to 24 hpi.

In conclusion, this reproducible *in vitro* human macrophage infection model is applicable to both *S*. Typhimurium and *S*. Enteritidis serovars. The platform enables comparative studies of iNTS pathovariants and provides a reliable and reproducible framework for exploring host-pathogen interactions. While THP-1 cells may not capture the complexity of macrophage polarisation, they do represent a valuable tool for studying *Salmonella* infection phenotypes.

Our results clearly demonstrate that *S*. Typhimurium ST313 replicates and survives more effectively within human macrophages than the *S*. Typhimurium ST19 pathovariant. We have extended previous observations by showing that bloodstream-associated iNTS isolates of *S*. Enteritidis also exhibit robust intra-macrophage growth and survival.

We conclude that both the iNTS *S*. Typhimurium and *S*. Enteritidis lineages that evolved in Africa exhibit hyper-replication in human THP-1-derived macrophages. This robust intracellular growth phenotype provides a powerful tool for dissecting the molecular basis of iNTS pathogenesis and will support future functional genomic investigations.

### Limitations of the study

PMA-differentiated THP-1 macrophages, while highly reproducible, may not fully recapitulate the phenotypic diversity and plasticity of primary human macrophages.

## RESOURCE AVAILABILITY

### Lead Contact

Requests for information and resources should be directed to Natalia Cattelan

### Materials Availability

Unique reagents generated in this study are available from the lead contact with a completed materials transfer agreement.

### Data and Code Availability

All data reported in this paper will be shared by the lead contact upon request.

All original code has been deposited at Github and is publicly available at https://github.com/Marien-kaefer/Count-objects-within-objects.git.

Any additional information required to reanalyze the data reported in this paper is available from the lead contact upon request.

## ACKNOWLEDGEMENTS

We are very grateful to Paul Loughnane for his expert technical assistance. We also thank all the present and former members of the Hinton Lab for helpful and productive discussions during this project.

## AUTHORS’ CONTRIBUTIONS

Conceptualization S.H.P., G.A.B., J.C.D.H and N.C.; methodology, F.A., S.H.P., M.H, and N.C; investigation F.A., M.H. and N.C; writing-original draft F.A. and N.C; writing-review & editing F.A., M.H., S.H.P, G.A.B., J.C.D.H, and N.C; funding acquisition G.A.B and J.C.D.H.; and supervision N.C.

## SUPPLEMENTAL INFORMATION

**Amadeo_Supplementary.docx Figures S1–S3, Tables S1–S3, and supplemental references**

## FIGURE TITLES AND LEGENDS

## STAR★METHODS

### THP-1 cell culture

THP-1 cells were purchased from ATCC (American Type Culture Collection: TIB-202) and grown in ATCC-modified RPMI 1640 medium (Gibco) supplemented with 10% (v/v) foetal bovine serum (FBS; Gibco) at 37 °C in a humidified incubator, with 5% CO_2_, according to ATCC recommendations.

### Bacterial strains and growth conditions

*S*. Typhimurium strains 4/74 (ST19) ^46^, D23580 (ST313 Lineage 2) ^46^ and BKQZM9 (ST313 Lineage 3) ^8^ and *S*. Enteritidis global epidemic strains P125109 and A1636 and the Central/Eastern African (CEA) strains D7795 and CP255 were used in this study ^47^. Strains were grown in LB broth and LB broth with 0.3M NaCl (inducing SPI1 medium). All isolates were maintained as frozen stocks at −80°C.

### Experimental assessment of the production of macrophage-like cells

THP-1 cells were plated into 24-well tissue culture treated plates (Corning) at a density of 6×10^5^ cells/well in the presence of phorbol 12-myristiate-12 acetate (PMA, Sigma-Aldrich). Different PMA concentrations and incubation time combinations were explored: (Condition A) PMA 10 ng/mL for 24 h, (Condition B) PMA 10 ng/mL for 24 h followed by a recovery in fresh RPMI 1640 media for 24 h, (Condition C) PMA 10 ng/mL for 48 h or (Condition D) PMA 50 ng/mL for 48h. Production of macrophage-like cells was assessed via cell adherence and upregulation of macrophage surface markers through flow cytometry.

### Optimised production of macrophage-like cells from THP-1 cells

Our experiments led us to adopt Condition D for the optimised differentiation of THP-1 cells into M0 macrophages, using RPMI 1640 medium containing 50 ng/mL PMA for 48 h. Multiplicities of infection (MOI) of either 5 or 10 were used, as indicated.

### Macrophage polarisation into pro- and anti-inflammatory like macrophages

Macrophage cells produced via Conditions B, C and D were further polarised into (1) M1-like macrophages by incubation with 10 ng/mL LPS from *Salmonella* Typhimurium (Sigma-Aldrich) and 20 ng/mL IFN-γ (Gibco) for 16 h, and (2) M2-like macrophages by incubation with 40 ng/mL rIL-4 (Gibco) and 20 ng/mL rIL-13 (Gibco) for 72 h. MOCK controls were run in parallel of both conditions ^28,34^.

### Flow cytometry

After THP-1 differentiation into macrophages, or their further polarisation into M1- and M2-like macrophages, cells were washed twice with PBS without Ca^2+^ and Mg^2+^ (Gibco) and detached with Versene (Gibco). Macrophage surface markers expression was assessed via flow cytometry by staining the cells with anti CD11b (FITC, #130-110-552, Miltenyi Biotec), anti CD14 (APC, #130-110-520, Miltenyi Biotec), anti CD36 (APC-Vio770, #130-110-743, Miltenyi Biotec), anti CD86 (PerCP-Vio700, #130-116-267, Miltenyi Biotec), anti HLA-DR (VioGreen, #130-111-795, Miltenyi Biotec), anti CD163 (PE, #130-112-128, Miltenyi Biotec), anti CD206 (VioBlue, #130-127-809, Miltenyi Biotec) according to manufacturer’s instructions. Cells were stained as displayed in Supplementary Table 2. Unstained cells were used as controls. Data were obtained using a ZE5 Cell Analyser (BioRad) flow cytometer acquiring between 10^4^ to 10^5^ events per sample and analysed using the FlowJo software.

### RT-PCR

To assess M1 and M2 polarisation, total RNA was extracted by phenol-chloroform method using TRIzol, and the quality was checked using the BioAnalyser (Agilent). mRNA was reverse transcribed using SensiFAST™ cDNA Synthesis Kit (Meridian) and real-time PCRs (RT-PCR) were performed to detect IL1β, TNFα, HLA-DR, CD163, TGFβ, CCL-18 and CCL-22 using the SensiFAST™ SYBR Green Kit (Meridian). RPL37A and ACTB were used as reference genes for normalisation. The sequence of the primers is listed in Supplementary Table 3. The 2^−ΔΔCt^ method was used to calculate the fold change of the genes in treated vs control samples for each condition.

### ELISA

Cytokine secretion into the culture media of polarised cells was assessed via ELISA sandwich kits (Invitrogen) according to manufacturer instructions. Pro-inflammatory IL-6 (#88-7066-22) and TNFα (#88-7346-22) and anti-inflammatory IL-10 (#88-7346-22) and TGFβ1 (#88-8350-22) cytokine levels were measured using the CLARIOStar plate reader.

### Salmonella survival in Macrophages

*Salmonella* invasiveness and survival in macrophages was evaluated via Gentamicin protection assay ^48^ and CFU analysis at 1.5, 4, 8 and 24 hours post infection (hpi). THP-1 cells were differentiated into Mø macrophages with RPMI 1640 medium containing 50 ng/mL PMA for 48 h. Prior to infection, *Salmonella* isolates were grown in LB broth at 37°C in a shaking incubator at 220 rpm for 18 h, then sub-cultured to log phase (OD_600_ ≈ 0.9) with inSPI1 media at 37°C. Concentration of each strain when OD_600_ ≈ 0.9 was established via CFU assay in a separate set of experiments and used to perform infection experiments with multiplicity of infections (MOIs) of 5 and 10. All infection steps, unless specified otherwise, were performed at 37°C in a 5% CO_2_ incubator. Each bacteria strain was suspended in Hank’s Balanced Salt Solution (HBSS; Gibco) and used to infect the macrophages for 1h. Each MOIs was evaluated in triplicate for each time point. Cells were then washed twice with HBSS and 0.5 mL of RPMI 1640 containing 100 μg/mL Gentamicin were added to each well for 30 minutes to eliminate any remaining extracellular bacteria. The set of infected cells used to calculate the CFU at 1.5 hpi were washed twice with HBSS and lysed using 1 mL/well of 0.1% w/v Sodium Deoxycholate (DOC) in PBS for 10 minutes at RT. The released bacteria were serially diluted in PBS and plated on LB agar plates to perform the CFU analysis. In parallel, for those sets of infected cells intended to assess CFU at 4, 8 and 24 hpi, cells were washed twice with HBSS and 1 mL fresh complete media containing 5 μg/mL Gentamicin was added to each well to prevent extracellular growth during further incubation. 20 minutes before the end of each incubation time point media was replaced again with RPMI 1640 containing 100 μg/mL Gentamicin and after 20 minutes cells were lysed with 0.1 % w/v DOC to perform CFU assay as explained above.

### Cytotoxicity

To assess cytotoxicity and cell death during infections, THP-1 cells were seeded into 24-well plates at a density of 6×10^5^ cells/well, differentiated into macrophages with 50 ng/mL PMA for 48 h and infected as described above. At the end of the 100 μg/mL Gentamicin treatment (1.5 hpi), cells were washed twice with HBSS and 1 mL of fresh complete media containing 5 μg/mL Gentamicin and 20 nM SYTOX Green (Invitrogen) was added to the infected cells. The plate was placed in an IncuCyte S3 (Sartorius) in a 37°C 5% CO2 incubator and four phase-contrast and green fluorescence images were taken per well with the 10x objective at 2, 4, 8 and 24 hpi. In parallel with the infection, a set of uninfected cells were incubated with fresh media with 20 mM SYTOX Green to assess cell death unrelated to the infection and a set of uninfected cells were treated with 0.1% Triton X-100 to kill all the cells and used to obtain the “100% cell death” SYTOX Green count. Finally, a set of uninfected macrophages not receiving the SYTOX Green were used to calibrate the machine. The analysis of the images was performed using the machine integrated software. The percentage of cell death was calculated as follow:

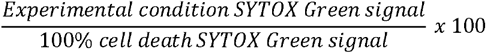

### Immunofluorescence analysis

Infection rates at MOI of 10 for each *Salmonella* isolate were assessed by immunofluorescence assays. THP-1 cells were seeded on sterile 13 mm diameter coverslips, differentiated into Mø macrophages with PMA 50 ng/mL for 48 h and infected with *Salmonella* as described above. After 1.5, 4, 8 and 24 hpi, cells were washed twice with PBS and fixed with PFA 4% for 20 minutes at RT. Cells were then washed twice with PBS and labelled with a *Salmonella* O antiserum factor 12 (BD Difco 227791), followed by an anti-rabbit 594 antibody (Invitrogen A11012), Alexa Fluor 488 Phalloidin (Invitrogen A12379) and DAPI (Thermo Scientific 62247) staining. Samples were mounted with ProLong Diamond antifade mounting media (Invitrogen P36965). Images were acquired on a Zeiss LSM 880 confocal microscope (Axio Examiner stand) using a Plan-Apochromat 63×/1.40 oil DIC objective (immersion oil, n = 1.518). Samples were imaged in line-sequential mode with three laser lines: 405 nm (diode) for DAPI, 488 nm (argon) for Alexa Fluor 488, and 561 nm (DPSS 561-10) for Alexa Fluor 594, with the corresponding main beam splitters (MBS-405 and MBS 488/561). Emission was collected on descanned PMT detectors with the following detection windows: 410–495 nm (DAPI), 498–587 nm (Alexa Fluor 488) and 587–733 nm (Alexa Fluor 594); the pinhole was set to ∼1 Airy unit (reported ∼1–2 AU across channels). Images were acquired as a z-stack with a z-step of 0.7 µm in stack/focus mode and line averaging of 2. Experiments were repeated independently three times. At least 100 cells per experiment were analysed.

### Image analysis

Image analysis was performed using Fiji (http://fiji.sc, ^49^) in combination with Cellpose (version 2.2.1, ^50^) and StarDist ^51^). Raw multichannel image stacks were first stitched in ZEN software and converted to 2D sum projections in Fiji. Mammalian cell segmentation was carried out using Cellpose (Cyto2 model) with optimized parameters for cell diameter (200-300) and probability thresholds (−3). Resulting instance masks were exported as PNG files and paired with the corresponding sum projection images.

Quantitative data were obtained from images analysed by means of Cellpose and StarDist macros run in Python and ImageJ respectively. Detailed explanation of image analysis and script can be found in https://github.com/Marien-kaefer/Count-objects-within-objects.git. Bacterial segmentation was performed on the bacterial fluorescence channel using StarDist with user-defined parameters (probability score/threshold: 0.35, overlap threshold: 0.6-0.7). The custom Fiji macro StarDist_for_child_objects_then_count_per_parent.ijm (available on Github) was used to associate segmented bacteria with Cellpose-derived cell regions of interest. For each cell, the number of unique bacterial objects within its boundary was quantified, and the results exported as CSV tables. Quality control was conducted by visual inspection of cell masks and bacterial segmentation outputs, with erroneous regions manually removed prior to analysis where necessary.

All analyses were run on a GPU-enabled workstation (HP Z6 G4, operating system: Windows 10 Enterprise 2016 LTSB Version 10.0.14393 Build 14393, GPU: NVIDIA Quadro RTX 4000, CUDA version: 10.0.130).

### Statistical analysis

All values in graphs are represented as mean ± standard deviation (SD), unless indicated otherwise in the figure legend. The statistical analysis was performed using the GraphPad Prism 9 software. The type of statistical test and the number of replicates included in each analysis are indicated in the figure captions.

