## supplementary for "Bloodstream-associated *Salmonella* Typhimurium and Enteritidis iNTS pathovariants hyper-replicate in human macrophages"

Supplementary material

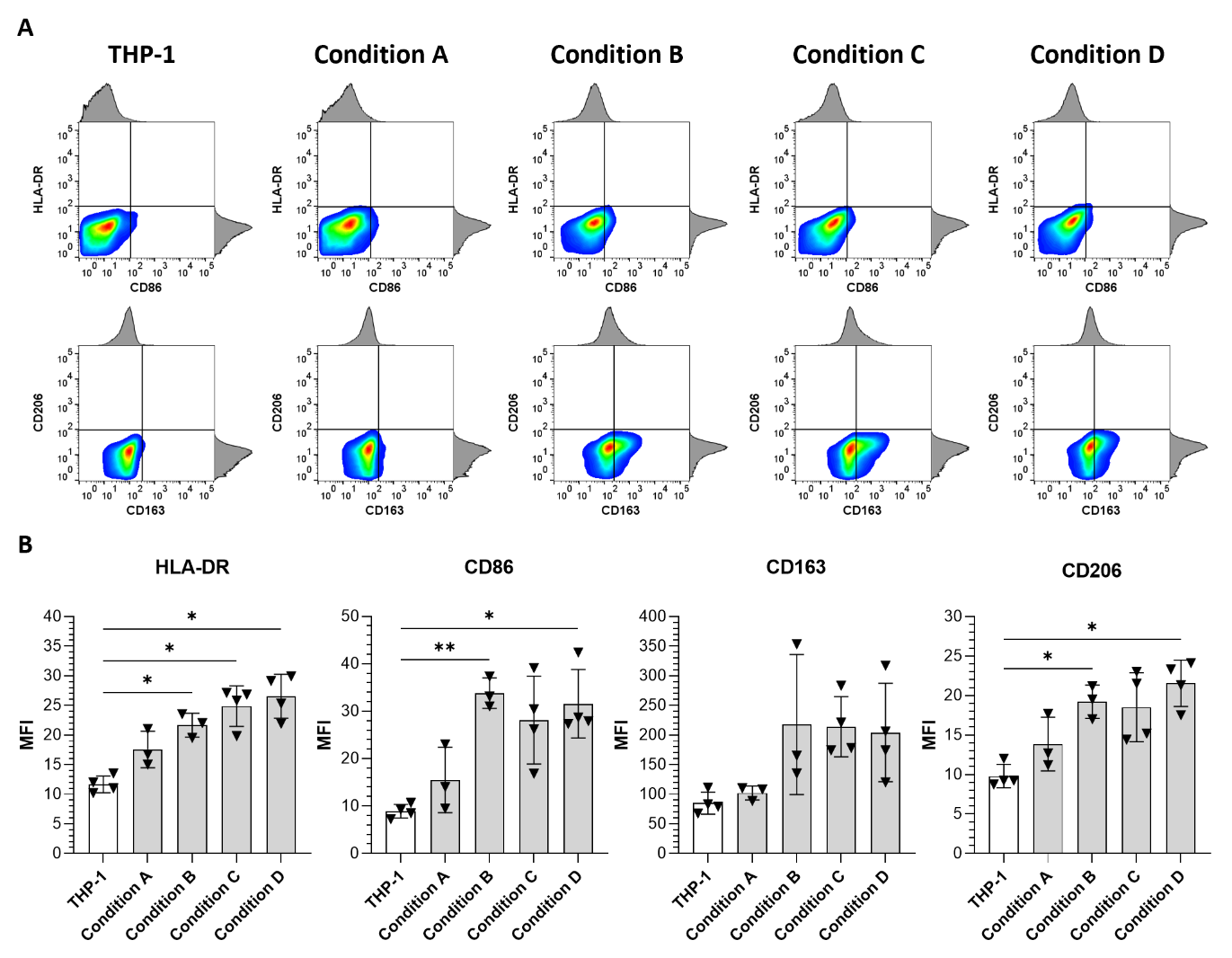

**Supplementary Figure 1|** **PMA does not activate THP-1 macrophages.** **(A)** Representative flow cytometric analysis of THP-1 cells versus differentiated THP-1 cells with Condition A (PMA 10 ng/mL for 24 h), Condition B (PMA 10 ng/mL for 24 h followed by a 24 h recovery in fresh media), Condition C (PMA 10 ng/mL for 48 h) and Condition D (PMA 50 ng/mL for 48 h) stained using anti CD86, anti HLA-DR or anti CD163 and anti CD206. **(B)** Median fluorescence intensity (MFI) of CD86, HLA-DR, CD163 and CD206-stained cells. Data are displayed as mean ± SD from n ≥ 3. Statistical analyses performed using the Brown‑Forsythe and Welch ANOVA test and the Dunnet T3 multiple comparison post hoc test. * p < 0.05, ** p < 0.01.

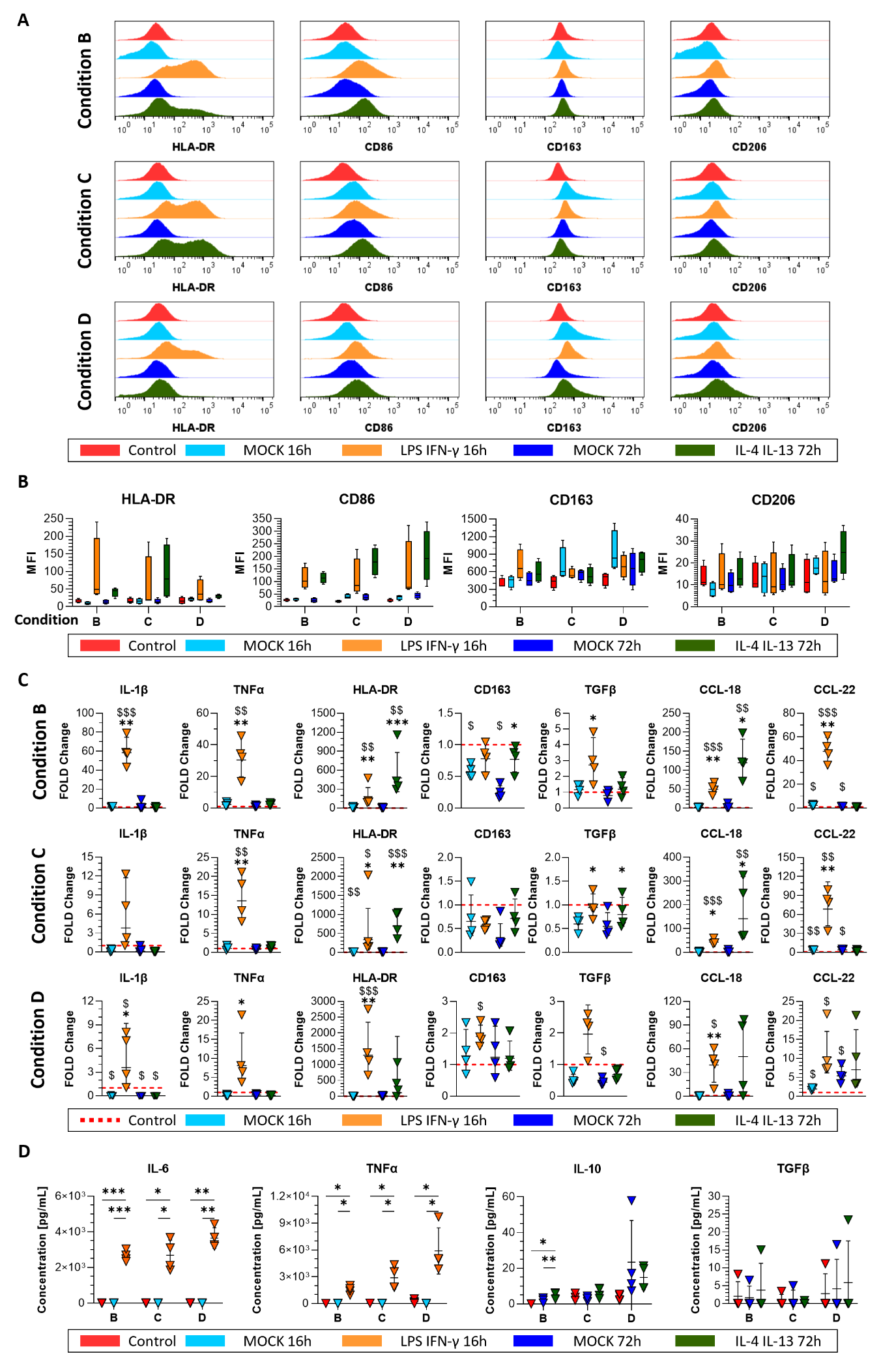

**Supplementary Figure 2|** **THP-1 derived macrophages can be activated to pro-inflammatory M1 macrophages. (A)** Representative flow cytometric panel of M1 (CD86 and HLA-DR) and M2 (CD163 and CD206) typical markers in macrophages obtained with Conditions B, C and D (Control, red) versus macrophage treated with LPS and IFN-γ for 16h (orange) or with IL-4 and IL-13 (light green) and the respective MOCKs (blue and dark green respectively). **(B)** Median fluorescence intensity (MFI) of CD86, HLA-DR, CD163 and CD206- stained cells. Data are displayed as mean ± SD from n ≥ 3. Statistical analyses performed using the Brown‑Forsythe and Welch ANOVA test and the Dunnet T3 multiple comparison post hoc test. **(C)** mRNA levels of M1 (IL-1β, TNFα and HLA-DR) and M2 (CD163, TGFβ, CCL‑18 and CCL‑22) associated genes detected via RT-PCR represented and displayed as 2-fold change. For each condition (B, C and D) data were normalised against the respective macrophage control (Control, red dashed line). Data are displayed as geometric mean ± geometric SD from n = 4. Statistical analyses performed on the DCt using the one-way ANOVA test and the Šidák multiple comparison post hoc test. $ refers to comparisons with each condition control while * with the respective mock. $ p < 0.05 v Control, $$ p < 0.01 v Control, $$$ p < 0.001 v Control, * p < 0.05 v mock, ** p < 0.01 v mock, *** p < 0.001 v mock. **(D)** Supernatant levels of pro- (IL-6, TNFα) and anti-inflammatory (IL-10, TGFβ) cytokines released by the THP-1 cells differentiated into macrophages following conditions B, C and D (Controls), the cells treated with LPS and IFN (orange), treated with IL4 and IL13 (dark green) and the respective MOCKs (blue and light blue). Concentrations are expressed as pg/mL. Data are displayed as mean ± SD from n = 3. Statistical analysis performed using a two-way ANOVA and the Tukey multiple comparison post hoc test. * p < 0.05, ** p < 0.01, *** p < 0.001.

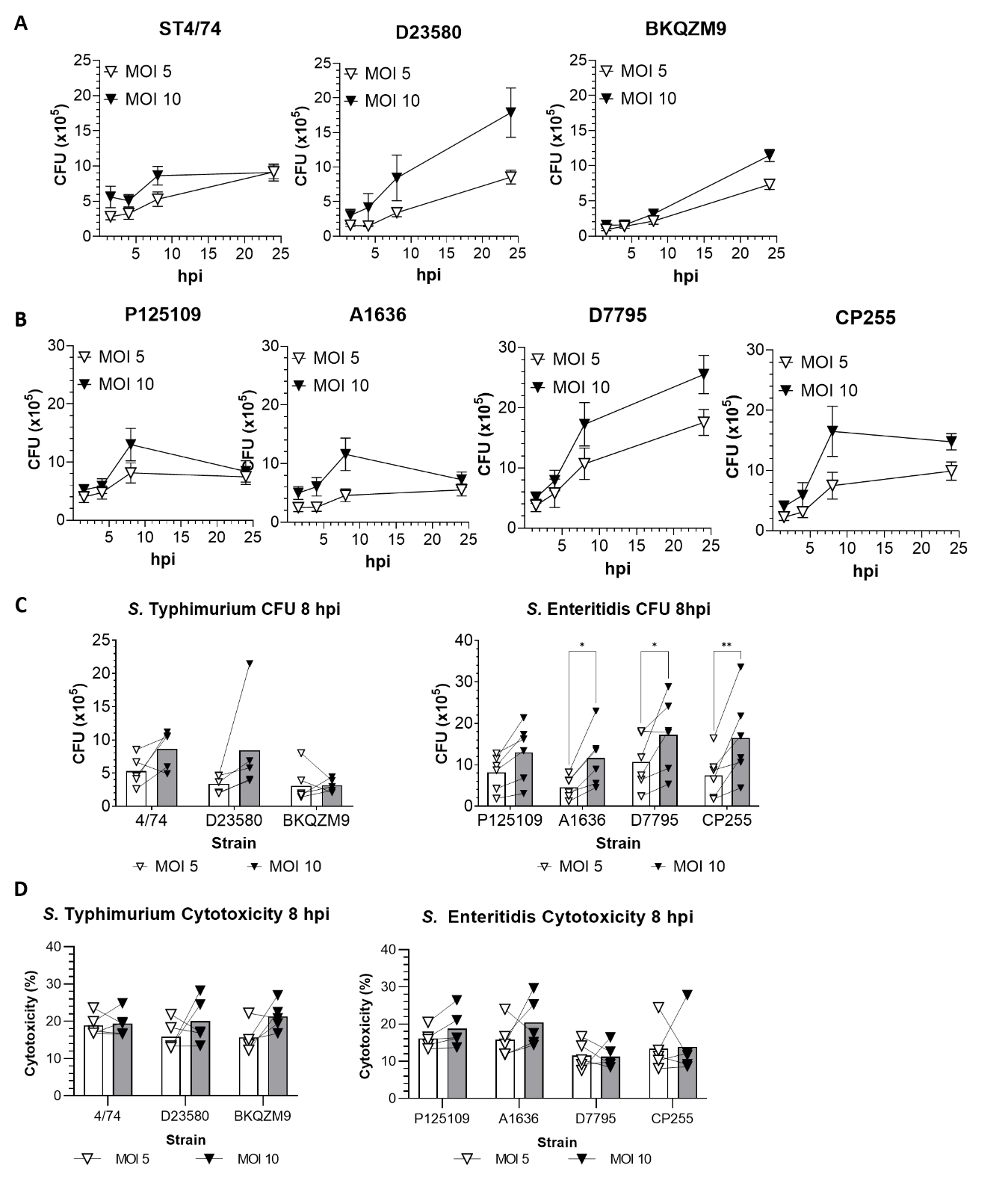

**Supplementary Figure 3|** **Varying the MOI did not alter bacterial behaviour inside macrophages.** THP-1 cells infected at a MOI of 5 and 10 with **(A)** *S.* Typhimurium ST19 (4/74) and ST313 L2 (D23580) and L3 (BKQZM9) and **(B)** *S.* Enteritidis GEC (P125109 and A1636) and CEAC (D7795 and CP255). **(A-B)** Intracellular bacteria were enumerated via CFU at 1.5, 4, 8 and 24 hpi. Data are displayed as mean ± SD from n = 6. **(C)** CFU at 8 hpi to compare the difference between MOI of 5 and MOI of 10 in *S.* Typhimurium and *S.* Enteritidis. Data are displayed as mean with experimental replicates connected by a line (n = 5). Statistical analysis performed using ratio paired t-tests. * p < 0.05, ** p < 0.01, *** p < 0.001. **(D)** Cytotoxicity (%) induced by *S.* Typhimurium and *S.* Enteritidis at 8 hpi comparing MOI of 5 and MOI of 10. Data are displayed as mean with experimental replicates connected by a line (n = 5).

**Supplementary Table 1:** THP-1 differentiation protocols used in different *Salmonella* articles.

|  | **PMA concentration (ng/ml)** | **Days of differentiation** | **Rest days** | **MOI infection** | **Reference** |
| --- | --- | --- | --- | --- | --- |
|  | Not specified | 3 | 1 | 20 | Van Puyvelde et al. ^1^ |
|  | 6.16 | 1 | No | 10 | Ramachandran et al. ^2^ |
|  | 20 | 1 | No | 10 | Valle et al. ^3^ |
|  | 20 | 3 | no | 40 | Starr et al. ^4^ |
|  | 20 | 5 | No | 10 | Hurley et al. ^5^ |
|  | 25 | 1 | No | 30 | Xu et al. ^6^ |
|  | 25 | 3 | No | 10 | Ingle et al. ^7^ |
|  | 50 | 2 | 4 | 5 | Winter et al. ^8^ |
|  | 50 | 3 | 2 | 30 | Fisch et al. ^9^ |
|  | 50 | 2 | No | 100 | Spiegelhauer et al. ^10^ |
|  | 50 | 1 | 3 | Not indicated | Luk et al. ^11^ |
| 61.6 | | 2 | No | 10 | Crouse et al. ^12^ |
|  | 61.6 | 1 | No | 50 | Knuff-Janzen et al. ^13^ |
|  | 61.6 | 2 | No | 10 | Baldassarre et al. ^14^ |
|  | 100 | 1 | No | 5 | Spano et al. ^15^ |
|  | 100 | 2 | 1 | Varies per mutant | Yeung et al. ^16^ |
|  | 100 | 2 | 1 | 15 | Mylona et al. ^17^ |
|  | 100 | 1 | 1 | 100 | Ganguli et al. ^18^ |
|  | 123 | 1 | No | 20 | Naseer et al. ^19^ |
|  | 200 | 1 | 1 | 50 | Gram et al. ^20^ |

**Supplementary Table 2:** Cell markers used to identify differentiation.

| Cell type | Markers | | |
| --- | --- | --- | --- |
| Mø | CD11b FITC | CD14 APC | CD36 APC-Vio770 |
| M1 | CD86 PerCP-Vio700 | HLA-DR VioGreen |  |
| M2 | CD163 PE | CD206 VioBlue |  |

**Supplementary Table 3:** Primers used for the real time PCR.

| Primer | Sequence |
| --- | --- |
| IL-1β_Fw | GCCCTAAACAGATGAAGTGCTC |
| IL-1β_Rv | GAGATTCGTAGCTGGATGCC |
| TNFα_Fw | CTGCACTTTGGAGTGATCGG |
| TNFα_Rv | TCAGCTTGAGGGTTTGCTAC |
| HLA-DR_Fw | CATAAGTGGAGTCCCTGTGCTA |
| HLA-DR_Rv | TCAGGATTCAGATAGAACTCGGC |
| CD163_Fw | TCGTCGCATTATTCTTCTTGACTAA |
| CD163_Rv | GGTGGACTAAGTTCTCTCCTCTTGA |
| TGFβ_Fw | GGAAATTGAGGGCTTTCGCC |
| TGFβ_Rv | CCGGTAGTGAACCCGTTGAT |
| CCL-18_Fw | AAGCCAGGTGTCATCCTCCTAAC |
| CCL-18_Rv | TGATGTATTTCTGGACCCACTTCTT |
| CCL-22_Fw | CACTTCTACTGGACCTCAGAC |
| CCL-22_Rv | AGTAGGCTCTTCATTGGCTCA |
| ACTB_Fw | TCCACGAAACTACCTTCAACTC |
| ACTB_Rv | CAGTGATCTCCTTCTGCATCC |
| RPL37A_Fw | ATTGAAATCAGCCAGCACGC |
| RPL37A_Rv | AGGAACCACAGTGCCAGATCC |
